# Development of PRPK Directed Phthalimides

**DOI:** 10.1101/2021.11.30.469594

**Authors:** Hyuk-Soo Seo, Takashi Mizutani, Teru Hideshima, Nicholas E. Vangos, Tinghu Zhang, Kenneth C. Anderson, Nathanael S. Gray, Sirano Dhe-Paganon

**Affiliations:** Department of Cancer Biology, Dana-Farber Cancer Institute, Boston, MA 02215; Department of Biological Chemistry & Molecular Pharmacology, Harvard Medical School, Boston, MA 02115; Jerome Lipper Multiple Myeloma Center, Department of Medical Oncology, Dana-Farber Cancer Institute, Harvard Medical School, Boston, MA 02215; Discovery and Preclinical Research Division, Taiho Pharmaceutical Co. Ltd., Tsukuba, Ibaraki 300-2611, Japan

**Keywords:** Immunomodulatory drugs, IMiDs, PRPK, TP53RK, TPRKB, crystal structures

## Abstract

Immunomodulatory drugs (IMiDs) thalidomide, lenalidomide, and pomalidomide (Pom) bind to cereblon (CRBN) and trigger proteasomal degradation of neo-substrates IKZF1/3 leading to multiple myeloma (MM) cell apoptosis. Pomalidomide (Pom) also binds albeit weakly to p53-related protein kinase (PRPK, aka TP53RK), an understudied kinase reported to be associated with poor prognosis in MM patients. Here, we developed a series of IMiDs based on Pom and conducted a structure-activity relationship (SAR) study to identify a potent and selective PRPK binder. Structural analysis showed that IMiDs bind PRPK in a fundamentally different way from CRBN, and suggested specific derivatization to improve affinity. We employed a structure-guided strategy to develop compound TXM-02-118, which exhibited nanomolar affinityfor PRPK in binding assays, and showed high selectivity for PRPK over CRBN. Overall, the work represents an initial effort to develop tool compounds for studying PRPK. Moreover, it illustrates how a single class of molecules can use different recognition elements to bind diverse targets using fundamentally different binding poses. This has broad implications for chemical probe and lead compound selectivity profiling, and argues for more wide-spread use of global proteomics or similar methodologies.

## Introduction

Immunomodulatory drugs (IMiDs), such as thalidomide, lenalidomide, and pomalidomide (Pom), are a class of FDA-approved drugs for treatment of erythema nodosum leprosum, myelodysplasia, and multiple myeloma (MM). Over the last decade, these agents have been of increasing interest given their unique mechanism of action. Unlike most other small molecule drugs that bind and modulate the activity of a given target, IMiDs mediate ternary complex formation between Cereblon (CRBN), a substrate receptor of the cullin-RING ubiquitin ligase (CRL) complex (Ito et al., 2010), and a neo-substrate, a cellular protein that is not recognized by CRBN in the absence of IMiDs. The best documented neo-substrates in this context are transcription factors IKZF1 and IKZF3 (Kronke et al., 2014; Lu et al., 2014). The functional outcome of IMiD-mediated complex formation between CRBN and neo-substrates, is neo-substrate polyubuiquitination and subsequent proteasomal degradation (Kronke et al., 2014; Lu et al., 2014). Thus, IMiDs belong to a growing number of molecules that induce targeted protein degradation and are often referred to as degrader molecules (Kostic and Jones, 2020; Wu et al., 2020).

More recently, knockdown studies have shown that Pom retained significant cellular potency in the absence of CRBN, suggesting that some Pom activity may be due to off-target effects, and proteomic analysis implicated p53-related protein kinase (PRPK, aka TP53RK) as a possible off-target (Hideshima et al., 2017). PRPK represents an understudied kinase as its pathophysiological role remains largely unknown. The limited amount of current literature suggests that PRPK is an atypical kinase that serves as an accessory subunit of the KEOPS complex, a large macromolecular assembly that catalyzes threonyl-carbamoyl-adenosinylation of tRNA (Perrochia et al., 2013). Unlike the catalytic subunit, called OSGEP, PRPK’s role in this complex is unclear, although a possibility that PRPK acts as an ATPase switch that induces a conformational change required for product release has been proposed (Mao et al., 2008; Zhang et al., 2015). Mutations in KEOPS-complex genes, including PRPK, have recently been associated with nephrotic syndrome with primary microcephaly (Braun et al., 2017; Wang et al., 2018) and Galloway-Mowat syndrome (Domingo-Gallego et al., 2019; Hyun et al., 2018). Independent of the KEOPS complex, PRPK has also been functionally implicated in telomere maintenance, p53 regulation, colorectal cancer metastasis, and skin cancer (Goswami et al., 2019a, 2019b; Hideshima et al., 2017; Roh et al., 2018; Zykova et al., 2018, 2017). Additionally, Depmap analysis suggests that PRPK has oncogenic properties, with this kinase being selectively essential for the growth of numerous cell types, including hematopoietic, breast and ovarian cancer lines (Ghandi et al., 2019). And increased PRPK levels confer poor prognosis in multiple myeloma (MM) patients and genetic knockdown of PRPK resulted in increased MM cell line apoptosis (Hideshima et al., 2017). Taken together, this body of evidence suggests that PRPK plays important regulatory roles, and that targeting and inhibiting PRPK may offer therapeutic opportunities. However, agents that selectively target PRPK have not been previously described. Here, we solved the structure of Pom bound to PRPK, revealing that CRBN and PRPK engage the IMiD scaffold in fundamentally different ways. We exploited this difference to develop a series of compounds based on the IMiD scaffold. The resulting lead compound, TXM-02-118, exhibited nanomolar affinity against PRPK in biochemical assays and no binding to CRBN. Therefore, TXM-02-118 represents the first IMiD-based compound that does not engage CRBN while potently binding to PRPK.

## Results and Discussion

### Crystal structure of human PRPK bound to Pom

We used recombinantly over-expressed full-length human PRPK/TPRKB complex for biochemical, biophysical and structural characterization of Pom binding. PRPK was unstable without its TPRKB subunit (p53-related protein kinase binding protein). The PRPK/TPRKB complex showed affinity of approximately 3 μM with Pom using ITC, ADP-Glo and TR-FRET (**Figure 1a**). The closely related IMiD compound Lenalidomide did not bind significantly to PRPK under the same conditions. To further characterize this biding, we co-crystallized the PRPK/TPRKB heterodimer with Pom and obtained crystals that diffracted to 2.3 Å resolution. This liganded structure was solved via molecular replacement with search models based on the previously reported bacterial structures and refined to Rwork and Rfree values of 0.17 and 0.22, respectively, and with data collection and refinement statistics summarized in **Table 2**. We observed that both PRPK and TPRKB assumed conformations similar to orthologous structures available in the PDB database (3ENP and 4WW5) with RMSD of 1.38 and 0.61 Å, respectively. The accessory TPRKB subunit was bound via the β-lobe of the catalytic PRPK subunit, interacting with the α-C helix and thus may have access and potentially may be a regulator of the catalytic machinery of PRPK (**Figure 2A**) (Modi and Dunbrack, 2019). Consistent with prokaryotic structures, the active site of human PRPK contains canonical positioning of ATP-binding and catalytic residues including the hinge region (Glu115-Ser119), glycine-rich aka p-loop (Gly42-Ala45), and a DFG loop (Asp183-Gly185) (**Figure 2B**). As with previous structures, the short, seven-residue activation loop was absent of secondary structure, well-packed against the core, and contained two hydroxyl-containing residues that were not significantly solvent exposed. These preliminary observations support previous suggestions that PRPK may not be a typical substrate- or peptide-processing protein kinase enzyme, but rather a conformational-change motor that could be required for threonyl-carbamoyl-adenosinylation of tRNA(Mao et al., 2008; Zhang et al., 2015).

**Figure 1.**
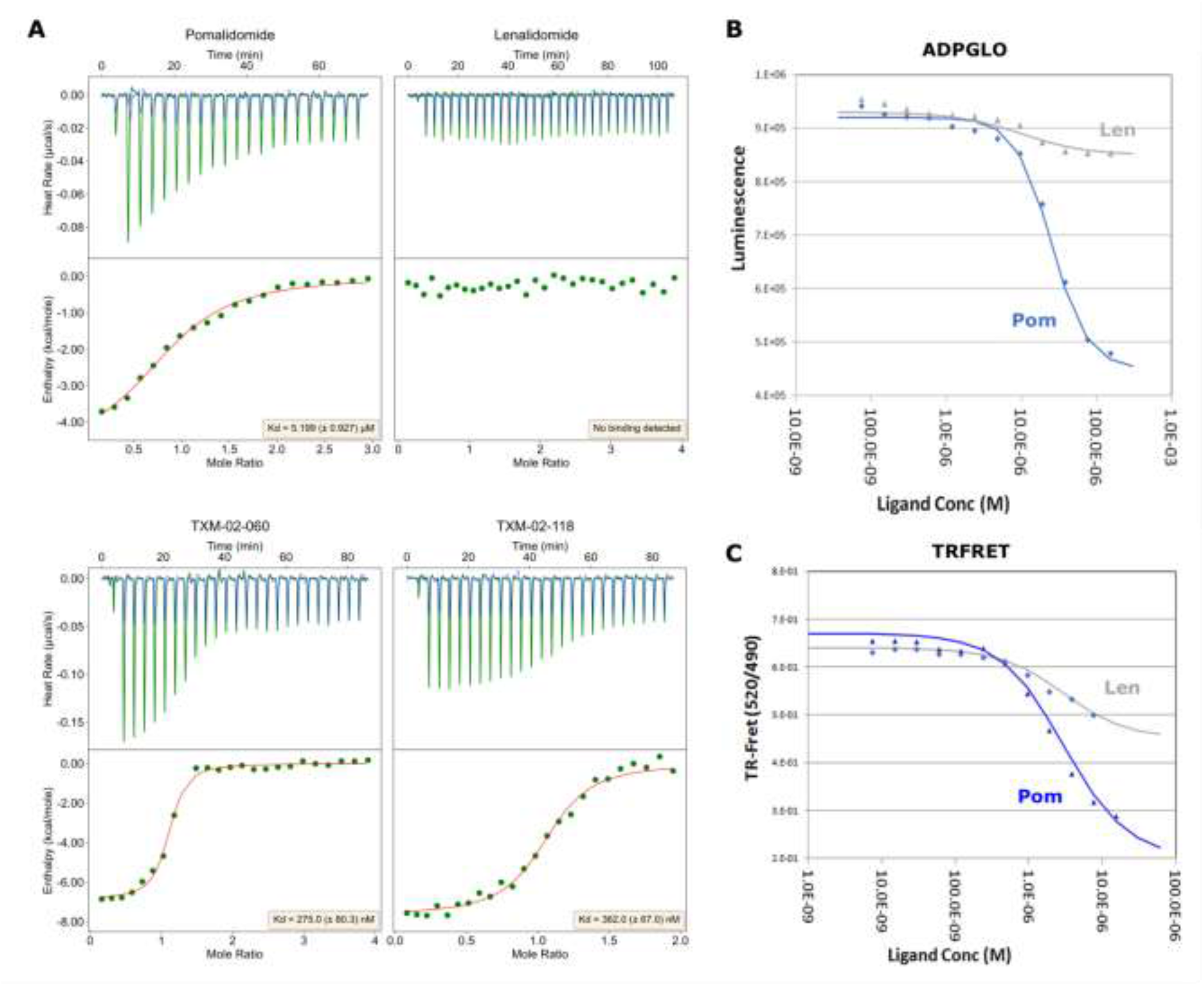
Biophysical characterization. (A) Isothermal titration calorimetry (ITC) measurements showing the binding of recombinant human PRPK/PRPKB complex to ligands in a 1:1 stoichiometry with binding affinity (dissociation constant, Kd) of ∼5.20, 0.27, and 0.36 uM for Pom, TXM-02-060, and TXM-02-118, respectively. ITC using Lenalidomide showed no heat of interaction. Data are representative of n = 3 independent ITC experiments with similar results. (B) A 12-dose serial-dilution response by ligands in the ADP-Glo assay measuring ADP production by PRPK complex to determine IC50s. Results of Pom and Len showed that the former but not the latter inhibited ATPase activity. (C) Direct biophysical binding assay was conducted using the TRFRET assay, showing that Pom bound with an IC50 of approximately 1uM while Len bound with an affinity that was approximately one order of magnitude lower.

**Figure 2.**
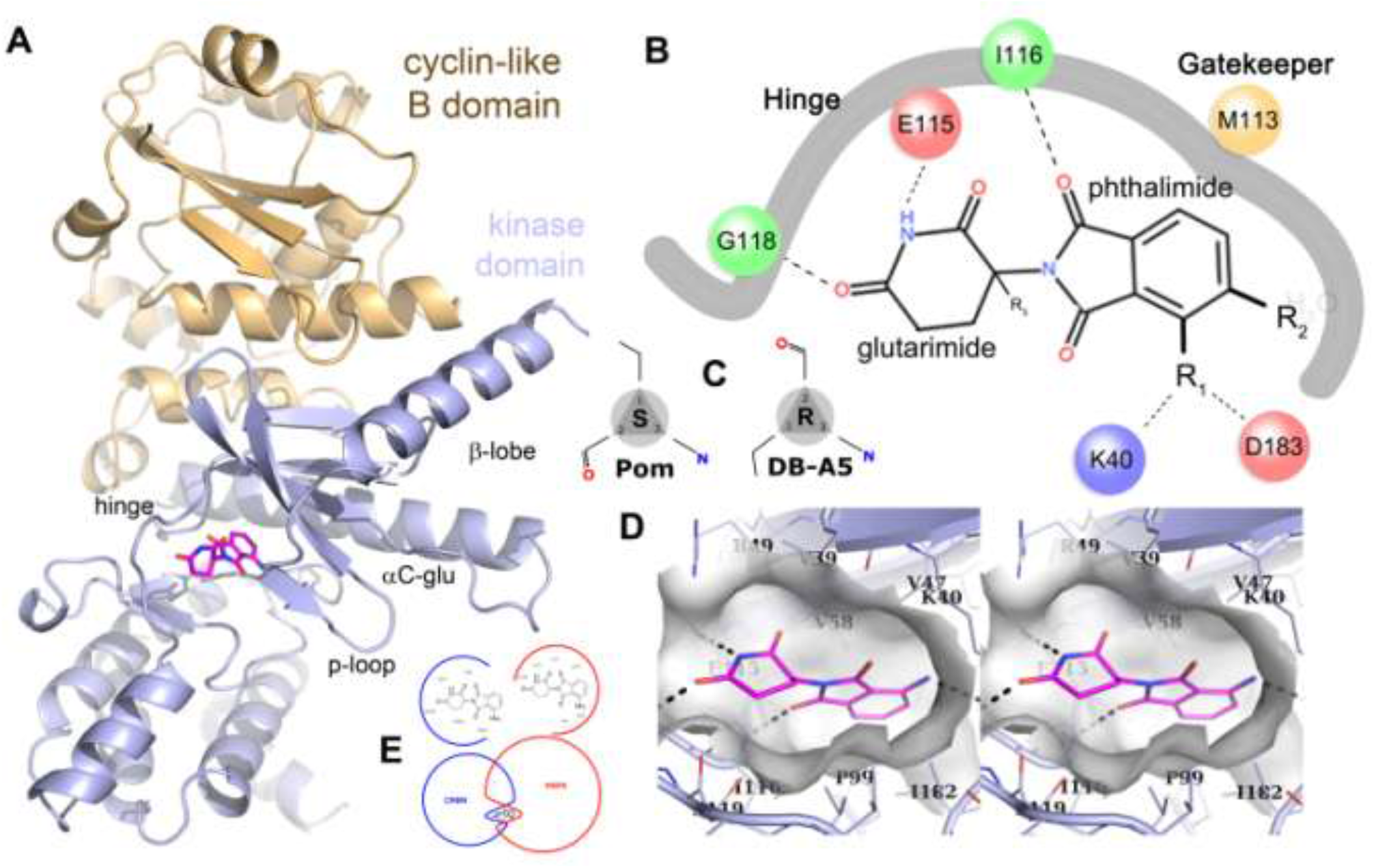
Structures of PRPK/PRPKB in complex with Pom. (A) Ribbon representation of PRPK, oriented in the standard protein kinase view, with Pom in stick format. (B) Ligand interaction diagram of the PRPK/Pom co-structure showing hydrogen bonds as dashed lines and the solid line representing boundaries within the active site that are close to the kinase backbone or towards deep, buried regions of the protein core. In contrast, R vectors that are either solvent exposed or point towards interaction-prone pockets within the active site represent potential opportunities for chemistry. (C) Fischer projections showing Pom in the S and DB-A5 in the R configurations, respectively. (D) Stereo view of PRPK (light blue) bound to Pom (magenta). Hydrogen bonds are indicated by dashed lines. (E) Ligand interaction diagram and slicing of the binding of Pom with CRBN and PRPK. Surface calculations show that these receptors are competitive with respect to each other for binding to Pom.

Due to the high resolution of the dataset, we were able to model Pom molecule into extra density located within the canonical ATP-binding site. We observed that the phthalimide (aka, isoindole) feature of Pom was buried in the canonical type I pocket near the gatekeeper methionine residue, and the glutarimid was partially solvent exposed (**Figure 2 and Figure 3A**). Pom formed a total of four well-defined hydrogen bonds, two by the isoindole and two by the glutarimid. Notably, the PRPK hinge region, including residues Glu115, Ile116, and Gly118, mediated three of the four hydrogen bonds, two of which were via backbone and one via the epsilon carboxyl side chain of Glu115. A water molecule was buried in the active site at the back-pocket near the gatekeeper, engaging the DFG segment and alphaC-Glu (Glu84). And the aliphatic ethylene portion of the glutarimid was solvent exposed. This binding pose suggested opportunities for chemical modifications of Pom to improve affinity.

**Figure 3.**
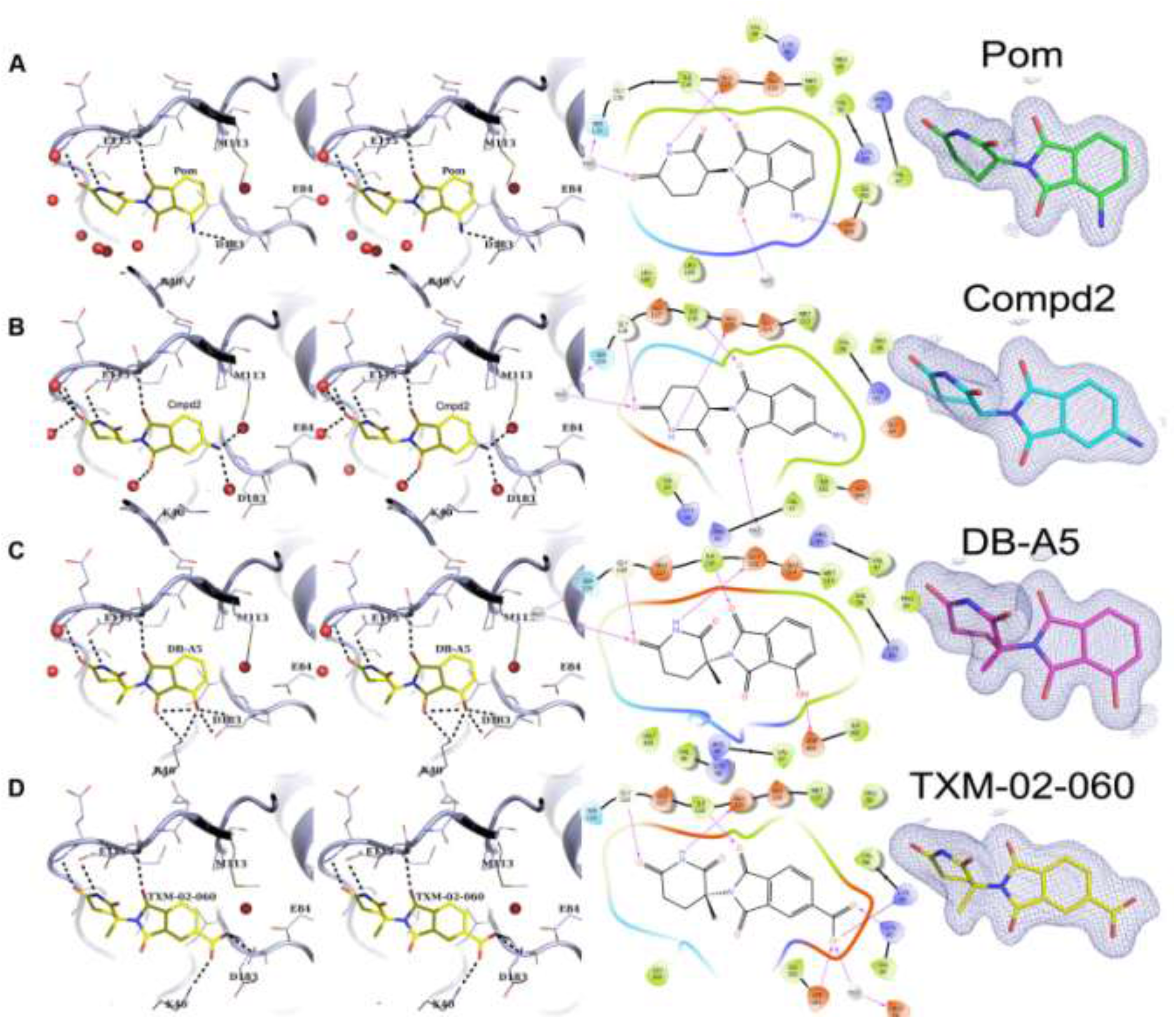
Structures of PRPK in complex with analogs. (A-D) complex structures of Pom, Compd2, DB-A5, and TXM-02-060. (Left) Stereoscopic representation in ribbon format of the active site in complex with analogs, showing sidechains as lines, hydrogen bonds as dashed lines, and waters as red spheres. (Middle) Ligand interaction diagrams of analog complex structures, showing hydrogen bonds as arrows. (Right) 2Fo-Fc map electron density map of ligands contoured at one sigma.

Pom contains a chiral center, with the R-configuration representing the form that is competent to bind to CRBN. However, the individual enantiomers racemize under normal physiological conditions due to the acidic hydrogen at the chiral center, and that both S- and R-IMiDs are most likely present *in vivo*. We noted that PRPK bound to the S configuration of Pom, but as this portion of the molecule was partially solvent exposed, there were no obvious restraints imposed by the protein on this configuration (**Figure 2 and Figure 3A**). While the overall quality of electron density for Pom was excellent, the density of the aliphatic ethylene portion of the glutarimid, which was solvent exposed, was relatively extended, consistent with the possibility that PRPK could bind either configuration.

In comparing the structures of PRPK bound to Pom and CRBN bound to Pom, we identified several key differences between how PRPK and CRBN engage Pom. Most notably, the glutarimid of Pom is buried when bound to CRBN, yet solvent exposed in complex with PRPK (**Figure 2E**). Therefore, our structure suggested that development of PRPK-selective IMiD-based compounds could be achieved through glutarimid-focused chemical derivatization and modifications, which are not tolerated by CRBN.

After characterizing Pom, we next examined a commercially-available Pom isomer, Compound2. Compound2 showed decreased PRPK binding affinity compared to Pom, and our structure of PRPK bound to Compound2 revealed that unlike Pom that engaged PRPK Asp183 and Lys40 via a hydrogen bond stemming from its amino group, Compound2’s amino group hydrogen bonded to two water molecules (**Figure 3B**). One water molecule was coordinated by the backbone amide of Lys182 and the carboxyl sidechain of GLu84 and sandwiched by van der Waals interaction by the sulfur atom of Met113 and the sidechains of Leu88 and Pro99 (**Figure 3B**). The other water molecule displaced the carboxyl sidechain of Asp183, flipping it into the solvent. This second water molecule also bonded with the amino group of Lys40. The primary amino group of Compound2 lost a hydrogen bond with the sidechain of Asp183 but gained two bonds with five waters. This interaction swap only slightly decreased overall observable affinity as measured by ITC.

### Structure-Guided Optimization of Pom Derivatives towards selective PRPK binding

The high-resolution structures of PRPK/PRPKB bound by Pom and Compound2 enabled rapid optimization of potency of IMiD-like derivatives. As shown in **Table 1**, we progressed towards compounds that displayed sub-μM K_D_ values for PRPK binding and showed no ability to bind CRBN.

**Table 1.** Evaluation of compounds against PRPK and CRBN. R1, R2, and R3 are shown represented in Figure 2. ITC-derived Kd values, in μM with standard error, of each compound for PRPK and CRBN are listed in the last two columns. Values labeled as >1000 did not show detectable heat of interaction.

| compounds | R1 | R2 | R3 | ITC/PRPK ($K_D$, μM) | ITC/CRBN ($K_D$, μM) |
| --- | --- | --- | --- | --- | --- |
| Pomalidomide | NH<sub>2</sub> | H | H | 5.2 ± 2.2 | 5.1 ± 2.8 |
| Compound2 | H | NH<sub>2</sub> | H | 6.1 ± 0.5 | 1.6 |
| TXM-01-027-1 | OH | H | H | 2.2 | NT |
| TXM-01-158 | OH | 4-pyrazole | H | 1.9 ± 1.7 | 0.54 |
| DB-A5 | OH | H | (*R*)-Me | 1.8 ± 0.4 | >1000 |
| DB-B5 | OH | H | (*S*)-Me | 27 | 19 |
| DB-A5-OMe | OMe | H | (*R*)-Me | >1000 | >1000 |
| TXM-02-071 | OH | 4-pyrazole | (*R*)-Me | >1000 | >1000 |
| TXM-02-118 | H | 5-(1,3,4-oxadiazole-2(3H)-thione) | (*R*)-Me | 0.36 ± 0.07 | >1000 |
| TXM-02-060 | H | CO<sub>2</sub>H | (*R*)-Me | 0.28 ± 0.08 | >1000 |

**Table 2.**
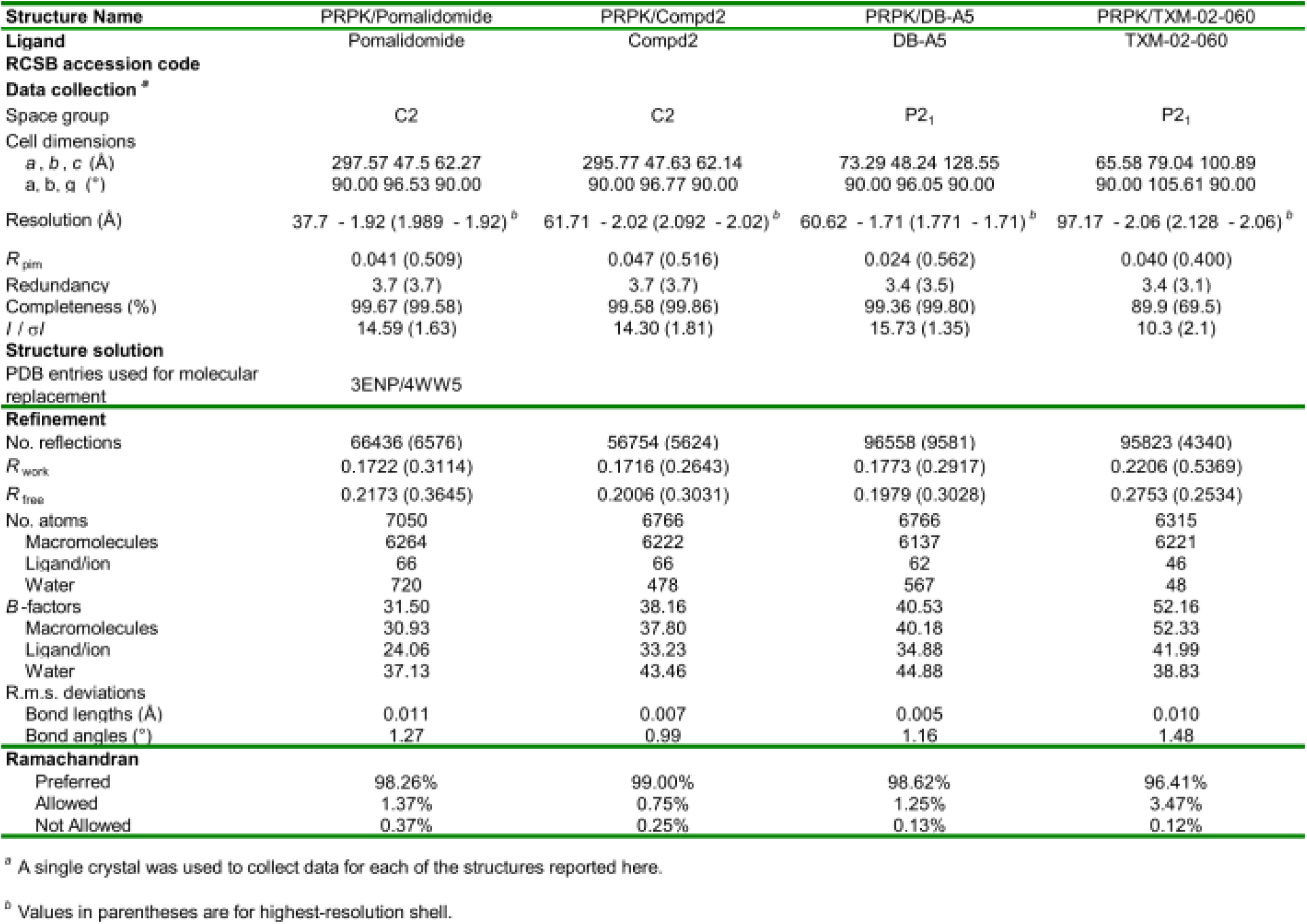
Crystallization conditions and data collection and refinement statistics.

We rationalized that the availability of a number of hydrogen bonding options at the R1 and R2 site could accommodate analogs with increased affinity, and we prepared TXM-01-027-1, which had a hydroxyl group at the R1 position. Indeed it showed improved potency, but only marginally as measured with ITC.

Introduction of 4-pyrazole at the R2 position (TXM-01-158) was tolerated but did not significantly improve potency. The lack of significant improvement when both R1 and R2 contain hydrogen binding moieties suggests competition between these moieties for interaction with the same protein residues, possibly Asp183. These preliminary SAR results suggested that the interaction of phthalimide derivatives with the PRPK was anchored by the hinge region and that the pocket formed by Lys40 and Asp183 and nearby water molecules represented an area that tolerates modifications, some of which could lead to significantly improved binding.

Previous work that reported PRPK as an off-target for Pom employed a probe that featured derivatization of Pom at the R3 position as an enrichment reagent (Hideshima et al., 2017), thus suggesting that the R3 position should be explored in the context of our SAR work described here. To explore this position and its contribution to PRPK affinity and specificity, we synthesized a series of R3-alkylated derivatives. Installing a methyl group at R3 of Pom did not improve PRPK binding but dramatically affected selectivity with respect to CRBN (**Table 1**). While the *R*stereoisomer (compound DB-A5) maintained single-digit micromolar affinity for PRPK, it completely abrogated binding to CRBN. On the other hand, the *S* stereoisomer (compound DB-B5) was selective for CRBN over PRPK. To explore this region, we determined a high resolution structure of PRPK bound to DB-A5 (Figure 3C). It showed a hydrogen bonding network similar to Pom, including the maintenance of all three hinge bonds and direct interaction of the R1 hydroxyl group with Asp183. Unlike Pom, DB-A5’s R1 hydroxyl group was able to recruit and bonded directly with the epsilon amino side chain of Lys40 (**Figure 3C**).

On the glutarimid side, the *R*-methyl group was pointed towards the bulk solvent (**Figure 3C**). To further explore the R3 position as a potential site for improved potency, we prepared a variety of R3 substituents, such as isopropyl or benzyl groups, but surprisingly none of these additional derivatives had appreciable affinity for PRPK (data not shown).

To improve PRPK potency while maintaining negative selectivity over CRBN, we retained the (*R*)-Me group at the R3 position and explored substituents at the R2 position. Surprisingly, the R1=OH and R2=4-pyrazole analog (TXM-02-071) was completely inactive. We hypothesized that the ‘anchor’ hydrogen bonding between a hydroxyl at R1 position and Lys40 on PRPK does not tolerate the coexistence of R2 and R3 substitutions. Therefore, we removed the hydroxyl group from R1 position in the presence of R3=(*R*)-Me, and added either R2=carboxyl (TXM-02-060) or R2=oxadiazolethione (TXM-02-118). Resulting compounds showed significantly improved PRPK binding while displaying no measurable affinity for CRBN. A structure of PRPK bound to TXM-02-060 was determined, showing that the carboxyl group formed ionic salt bonds with Lys40 as well as Lys60. Recruitment of Lys60 was associated with relaxation and opening-up of the active site, increasing the distance between the p-loop Ala43 backbone and the alpha-lobe Asp162 backbone by approximately 2.1 Angstroms. The former observation tangentially suggested that PRPK may be able to access multiple conformations of the active site that would be expected to be required for a normal phosphorylation-based catalytic reaction cycle, and lends further credence that PRPK is not a pseudokinase.

In summary, our SAR showed that PRPK’s active site lysine and aspartic residues (Lys40 and Asp183) were productively engaged by ligands that contain polar groups in the region defined by R1 or R2 substituent, culminating in compound TXM-02-118, which displayed the highest PRPK affinity. We noted the presence of a mutual exclusivity between R1 and R2 substituents. Moreover, a methyl group at the R3 position of Pom was well-tolerated by PRPK while completely abolishing CRBN binding. Finally, although additional R3 derivatives were not tolerated, we nevertheless believed that further exploration along this route will be productive.

Our structural analysis highlights that IMiD compounds can engage targets other than CRBN using fundamentally different binding poses. This likely stems from the presence of recognition-enabling features on both phthalimide section of the molecule that engages CRBN, and the glutariimide side that is critical for engaging PRPK. We exploited these key differences to optimize Pom into a molecule, TXM-02-118, that binds tightly to PRPK but has no affinity for CRBN. This study has several important implications, such as: (1) that TXM-02-118 represents a potential starting point for future optimization for developing chemical probes for studying PRPK biology, as this kinase remains poorly characterized; (2) that IMiDs may have a broader scope of targets beyond CRBN, including as ATP-competitive kinase inhibitors, and additional mechanisms of action beyond targeted degradation; and (3) that small molecule modulators of biological activity could use multiple different binding poses to engage different classes of targets, stressing the need to include proteome-wide assessment of selectivity as a standard step in chemical probe and lead candidates validation.

### Abbreviations

AcOH, acetic acid, Ac_2_O, acetic anhydride, CETSA, cellular thermal shift assay; DCM, dichloromethane; DIEA, N,N-diisopropylethylamine; DMF, dimethylformamide; DMSO, dimethyl sulfoxide; EtOAc. Ethyl acetate, KOAc, potassium acetate, MS, mass spectrometry; Na2CO3, sodium carbonate, NMR, nuclear magnetic resonance; NMP, N-methyl-2-pyrrolidone; TEA, triethylamine; TFA, trifluoroacetic acid; THF, tetrahydrofuran; TMS, tetramethylsilane; UPLC, ultra-performance liquid chromatography; UV, ultraviolet;

## Materials and Methods

### Cloning, protein expression and purification

N-terminally His-tagged full-length, bicistronic construct of human TP53RK and PRPK in the pET28PP vector was overexpressed in E. coli BL21 (DE3) in LB medium in the presence of 50 mg/ml of kanamycin. Cells were grown at 37°C to an OD of 0.8, cooled to 17°C, induced with 500 μM isopropyl-1-thio-D-galactopyranoside, incubated overnight at 17°C, collected by centrifugation, and stored at −80°C. Cell pellets were sonicated in buffer A (50 mM hepes 7.5, 300 mM NaCl, 10% glycerol, 10 mM Imidazole, and 3 mM BME) and the resulting lysate was centrifuged at 30,000 xg for 30 min. Ni-NTA beads (Qiagen) were mixed with lysate supernatant for 30 min and washed with buffer A. Beads were transferred to an FPLC-compatible column and the bound protein was washed with 15% buffer B (50 mM hepes 7.5, 300 mM NaCl, 10% glycerol, 300 mM Imidazole, and 3 mM BME) and eluted with 100% buffer B. HRV3C was added to the eluted protein and incubated at 4°C overnight. The sample was passed through a Superdex 200 10/300 column (GE healthcare) in a buffer containing 20 mM hepes7.5, 200 mM NaCl, 5% glycerol, and 1mM TCEP. Fractions were diluted and subjected to Mono-S ion-exchange chromatography and protein was eluted using a NaCl gradient. Fractions containing equimolar TP53RK and PRPK were pooled, concentrated, and frozen at - 80°C.

### Crystallization, data collection and structure determination

Samples of ∼400 μM protein and ∼600 μM ligand were crystallized in various precipitants by sitting- or hanging-drop vapor diffusion at 20°C. Crystals were transferred briefly into crystallization buffer containing 25% glycerol prior to flash-freezing in liquid nitrogen. Diffraction data from complex crystals were collected at beamline 24ID-E of the NE-CAT at the Advanced Photon Source at the Argonne National Laboratory. Data sets were integrated and scaled using XDS (Kabsch, 2010). Structures were solved by molecular replacement using the program Phaser (McCoy et al., 2007) and the search model PDB entry 3ENP and 4WW5. Iterative model building, ligand fitting, and refinement using Phenix (Liebschner et al., 2019) and Coot (Emsley et al., 2010) led to models with excellent statistics.

### Isothermal titration calorimetry (ITC) measurements

All calorimetric experiments were carried out using an Affinity ITC from TA Instruments (New Castle, DE) equipped with auto sampler in a buffer containing 20 mM HEPES, pH 7.5, 150 mM NaCl, 0.5 mM TCEP, and 2% DMSO at 25 °C. 20 μM protein solution in the calorimetric cell was titrated with 200 μM compound solution using 2 μL injection in 200 sec intervals using stirring speed at 125rpm. Resulting isotherm was fitted with a single site model to yield thermodynamic parameters of ΔH, ΔS, stoichiometry, and K_D_ using NanoAnalyze software (TA instruments).

### Differential Scanning Fluorimetry (DSF)

Experiments were carried out in a RTPCR 7500 Real-Time System (LifeTech) in 96 well plates, a total volume of 20 ul, and with an optimized SYPRO Orange dye concentration (5000x concentration in DMSO, Invitrogen). Compound dilutions in assay buffer (25 mM HEPES pH 7.5, 150 mM NaCl, 1 mM TCEP) were first prepared by an NT8 liquid handler (Formulatrix) and then addition of dye was performed also by the same liquid handler. Sealed plates were heated at 1°C/min from 25°C to 95°C with fluorescence readings every 0.5°C. Tm values were determined as the minimum of the first derivative of the recorded fluorescence intensity versus temperature plot.

### ADP-GLO assay

Inhibitor was titrated in protein sample containing 100uM NaOrthovanadate, and this was followed by a 30 minute incubation at 20°C, addition of ADPGLO reagent, a further incubation at 20°C for 40 mins, addition of KinaseDetection buffer, and a final incubation at 20°C for 40 mins. Luminescence was read using a Clariostar (BMG Labtech).

### Time-resolved fluorescence resonance energy transfer (TR-FRET)

Prepared was FITC-labelled (DB-08) at 500nM, his-tagged PRPK complex at 100 nM, terbium-coupled anti-his at 1nM (Invitrogen) and decreasing concentrations of compounds in 384-well microplates (Corning, 4514) in a buffer containing 50mM Tris pH 7.5, 150mM NaCl, 0.01% Tween20. Before measurements, reactions were incubated for 15 min at room temperature. After excitation of terbium (Tb) fluorescence at 337nm, emission at 490nm (Tb) and 520nm (FITC) were recorded with a 70 μs delay to reduce background fluorescence and the reaction was followed over 1 h by recording 60 technical replicates of each data point using a PHERAstar FS microplate reader (BMG Labtech). The TR-FRET signal of each data point was extracted by calculating the 520/490nm ratio. Data were analysed with MatPlotLib.

### Synthesis of compounds

Unless otherwise noted, reagents and solvents were used as received from commercial suppliers. All reactions were monitored using UPLC-MS: Waters Acquity UPLC/MS system (Waters Acquity QDa Detector, Waters Acquity PDA eλ Detector, Waters Acquity-I UPLC class Sample Manager-FTN, Waters Acquity-I UPLC class Binary Solvent Manager) using Acquity UPLC® BEH C18 column (2.1 × 50 mm, 1.7 μm particle size). solvent gradient = 85% A at 0 min,1%A at 1.7 min; solvent A = 0.1% formic acid in Water; solvent B = 0.1% formic acid in Acetonitrile; flow rate: 0.6mL/min. Purification of reaction product was carried out by flash chromatography using CombiFlash®Rf with Teledyne Isco RediSep® normal-phase silica flash columns; or Waters HPLC system using SunFireTM C18 column (19 × 100 mm, 5 μm particle size): solvent gradient 0% to 99% acetonitrile in H_2_O (0.035% TFA as additive); flow rate: 20 mL/min, or SunFireTM C18 column (30 × 250 mm, 5 μm particle size): solvent gradient 0% to 99% acetonitrile in H_2_O (0.035% TFA as additive); flow rate: 40 mL/min. 1H NMR spectra were obtained using 500MHz Bruker Avance III spectrometers. Chemical shifts are reported relative to deuterated methanol (δ = 3.31) or dimethyl sulfoxide (δ = 2.50) for 1H NMR. Spectra are given in ppm (δ) and as br = broad, s = singlet, d = doublet, t = triplet, q = quartet, m = multiplet and coupling constants (J) are reported in Hertz.

**Pomalidomide (Pom), 4-amino-2-(2,6-dioxopiperidin-3-yl)isoindoline-1,3-dione**, CAS 19171-19-8

**Compound2, 5-amino-2-(2,6-dioxopiperidin-3-yl)isoindoline-1,3-dione**, CAS 191732-76-0, purchased from Combi-Blocks

**TXM-01-027-1, 2-(2,6-dioxopiperidin-3-yl)-4-hydroxyisoindoline-1,3-dione**, CAS 5054-59-1, prepared as described in literature. (Bradner, James, et al, PCT Int. Appl., 2018148440, 16 Aug 2018)

**DB-A5, (*R*)-4-hydroxy-2-(3-methyl-2,6-dioxopiperidin-3-yl)isoindoline-1,3-dione**, prepared as described in literature. (Bradner, James, et al, PCT Int. Appl., 2018148440, 16 Aug 2018)

**DB-B5, (*S*)-4-hydroxy-2-(3-methyl-2,6-dioxopiperidin-3-yl)isoindoline-1,3-dione**, prepared as described in literature. (Bradner, James, et al, PCT Int. Appl., 2018148440, 16 Aug 2018)

### DB-A5-OMe, (*R*)-4-methoxy-2-(3-methyl-2,6-dioxopiperidin-3-yl)isoindoline-1,3-dione

The mixture of 4-methoxyisobenzofuran-1,3-dione (4.8 mg, 0.027 mmol), (*R*)-3-amino-3-methylpiperidine-2,6-dione hydrobromide (5.0 mg, 0.022 mmol) and potassium acetate (6.6 mg, 0.067 mmol) in AcOH (0.2 mL) was stirred at 90°C for 16 hrs. The reaction mixture was cooled and purified by reverse phase preparative HPLC (0-80% MeOH in water (0.035% TFA modifier)) to give the title compound (3.8 mg, 56%) as a colorless solid. 1H NMR (500 MHz, DMSO-d6) δ 10.98 (s, 1H), 7.81 (dd, J = 8.5, 7.3 Hz, 1H), 7.49 (d, J = 8.5 Hz, 1H), 7.39 (d, J = 7.3 Hz, 1H), 3.95 (s, 3H), 2.74 – 2.63 (m, 1H), 2.59 – 2.50 (m, 2H), 2.06 – 1.97 (m, 1H), 1.87 (s, 3H). LC-MS Mass RT=0.67 min, m/z: 303.4 [M+H]+.

### TXM-01-158

#### 4-bromo-3-methoxyphthalic acid

To the stirred solution of 3-bromo-2-methoxy-6-methylbenzoic acid (500 mg, 2.0 mmol) and KOH (572 mg, 10 mmol) in water (20 mL) was added potassium permanganase (967 mg, 6.1 mmol) at rt. After being stirred at 70°C for 3 days, NaHSO_3_ (637 mg, 6.1 mmol) was added at rt. The stirred mixture was carefully acidified with 6M HCl then the precipitate was collected by filtration and washed with water to give a colorless solid (426 mg, 76%). 1H NMR (500 MHz, DMSO-d6) δ 7.66 (d, J = 8.4 Hz, 1H), 7.55 (d, J = 8.4 Hz, 1H), 3.81 (s, 3H).

#### 5-bromo-2-(2,6-dioxopiperidin-3-yl)-4-methoxyisoindoline-1,3-dione

The suspension of 4-bromo-3-methoxyphthalic acid (200 mg, 0.73 mmol) in acetic anhydride (2 mL) was stirred at 130°C for 5 hrs then cooled down to rt. The stirred mixture was diluted with EtOAc and the precipitate was filtered off. The filtrate was evaporated with EtOAc and dried in vacuo to give crude compound as a dark purple solid. The mixture of crude 5-bromo-4-methoxyisobenzofuran-1,3-dione, 3-aminopiperidine-2,6-dione hydrochloride (132 mg, 0.80 mmol) and potassium acetate (215 mg, 2.2 mmol) in AcOH (5 mL) was stirred at 100°C for 16 hrs. The reaction mixture was cooled and partitioned between EtOAc and water. Organic layer was separated, washed with water then concentrated. The residue was purified by silica gel column chromatography (0 to 10% MeOH in DCM) to give the title compound (201 mg, 75%) as a dark orange solid. 1H NMR (500 MHz, DMSO-d6) δ 11.14 (s, 1H), 8.15 (d, J = 7.8 Hz, 1H), 7.57 (d, J = 7.8 Hz, 1H), 5.16 (dd, J = 12.9, 5.4 Hz, 1H), 4.08 (s, 3H), 2.89 (ddd, J = 17.2, 13.9, 5.5 Hz, 1H), 2.66 – 2.51 (m, 2H), 2.06 (m, 1H)..

#### 2-(2,6-dioxopiperidin-3-yl)-4-hydroxy-5-(1H-pyrazol-4-yl)isoindoline-1,3-dione (TXM-01-158)

The mixture of 5-bromo-2-(2,6-dioxopiperidin-3-yl)-4-methoxyisoindoline-1,3-dione (5.0 mg, 0.014 mmol), tert-butyl 4-(4,4,5,5-tetramethyl-1,3,2-dioxaborolan-2-yl)-1H-pyrazole-1-carboxylate (16 mg, 0.054 mmol), bis-(dibenzylideneacetone)-palladium (1.2 mg, 0.0014 mmol), 4,5-Bis(diphenylphosphino)-9,9-dimethylxanthene (1.6 mmol, 0.0028 mmol), dicyclohexylmethylamine (5.8 mL, 0.027 mmol) and 2M Na_2_CO_3_ (6.8 mL, 0.014 mmol) in 1,4-dioxane (0.2 mL) was stirred at 100°C for 16 hrs. The reaction mixture was purified by silica gel column chromatography (0-10% MeOH in DCM) to give crude 2-(2,6-dioxopiperidin-3-yl)-4-methoxy-5-(1H-pyrazol-4-yl)-isoindoline-1,3-dione as a pale brown gum. To the solution of the crude product in DCM (0.1 mL) was added borontribromide (1.0 M in DCM, 0.1 mL) at rt. After being stirred at rt for 16 hrs, reaction was quenched by the addition of water and the volatiles were removed under reduced pressure. The residue was purified by reverse phase preparative HPLC (0-80% MeOH in water (0.035% TFA modifier)) to give the title compound (1.1 mg, 24% in 2 steps) as a pale yellow solid. 1H NMR (500 MHz, DMSO-d6) δ 11.05 (s, 1H), 10.32 (s, 1H), 8.17 (brs, 2H), 7.98 (d, J = 7.6 Hz, 1H), 7.34 (d, J = 7.6 Hz, 1H), 5.04 (dd, J = 12.7, 5.4 Hz, 1H), 2.83 (ddd, J = 16.8, 13.5, 5.4 Hz, 1H), 2.57 – 2.45 (m, 2H), 2.04 – 1.94 (m, 1H). LC-MS Mass RT=0.50 min, m/z: 273.3 [M-H]-.

### TXM-02-071

#### (*R*)-5-bromo-4-methoxy-2-(3-methyl-2,6-dioxopiperidin-3-yl)isoindoline-1,3-dione

(*R*)-5-bromo-4-methoxy-2-(3-methyl-2,6-dioxopiperidin-3-yl)isoindoline-1,3-dione was synthesized as the same procedure as 5-bromo-2-(2,6-dioxopiperidin-3-yl)-4-methoxyisoindoline-1,3-dione, by using (*R*)-3-amino-3-methyl-piperidine-2,6-dione hydrobromide instead of 3-aminopiperidine-2,6-dione hydrochloride. 1H NMR (500 MHz, DMSO-d6) δ 11.02 (s, 1H), 8.11 (d, J = 7.8 Hz, 1H), 7.50 (d, J = 7.8 Hz, 1H), 4.03 (s, 3H), 2.68 (ddd, J = 18.3, 10.6, 5.4 Hz, 2H), 2.63 – 2.52 (m, 2H), 2.11 – 2.00 (m, 1H), 1.88 (s, 3H).

#### (*R*)-4-hydroxy-2-(3-methyl-2,6-dioxopiperidin-3-yl)-5-(1H-pyrazol-4-yl)isoindoline-1,3-dione (TXM-02-071)

(*R*)-4-hydroxy-2-(3-methyl-2,6-dioxopiperidin-3-yl)-5-(1H-pyrazol-4-yl)isoindoline-1,3-dione (TXM-02-071) was synthesized as the same procedure as 01-158, by using 4-(4,4,5,5-tetramethyl-1,3,2-dioxaborolan-2-yl)-1-((2-(trimethylsilyl)ethoxy)methyl)-1H-pyrazole instead of tert-butyl 4-(4,4,5,5-tetramethyl-1,3,2-dioxaborolan-2-yl)-1H-pyrazole-1-carboxylate. 1HNMR (500 MHz, DMSO-d6) δ 13.12 (brs, 1H), 11.02 (s, 1H), 10.27 (s, 1H), 8.37 – 8.08 (m, 2H), 8.03 (d, J = 7.6 Hz, 1H), 7.33 (d, J = 7.6 Hz, 1H), 2.76 – 2.65 (m, 1H), 2.62 – 2.52 (m, 2H), 2.10– 2.01 (m, 1H), 1.90 (s, 3H). LC-MS Mass RT=0.74 min, m/z: 354.9 [M+H]+.

### TXM-02-060

(*R*)-2-(3-methyl-2,6-dioxopiperidin-3-yl)-1,3-dioxoisoindoline-5-carboxylic acid (TXM-02-060) was synthesized as the same procedure as (*R*)-4-methoxy-2-(3-methyl-2,6-dioxopiperidin-3-yl)isoindoline-1,3-dione (DB-A5-OMe), by using 1,3-dioxo-1,3-dihydroisobenzofuran-5-carboxylic acid instead of 4-methoxyisobenzofuran-1,3-dione. 1H NMR (500 MHz, DMSO-d6) δ 13.77 (brs, 1H), 11.05 (s, 1H), 8.38 (dd, J = 7.8, 1.4 Hz, 1H), 8.21 (d, J = 1.4 Hz, 1H), 7.97 (d, J = 7.8 Hz, 1H), 2.74 – 2.64 (m, 1H), 2.64 – 2.53 (m, 2H), 2.07 (m, 1H), 1.91 (s, 3H). RT=0.68 min, m/z: 316.8 [M+H]+.

### TXM-02-118

#### tert-butyl (*R*)-2-(2-(3-methyl-2,6-dioxopiperidin-3-yl)-1,3-dioxoisoindoline-5-carbonyl)hydrazine-1-carboxylate

The mixture of (*R*)-2-(3-methyl-2,6-dioxopiperidin-3-yl)-1,3-dioxoisoindoline-5-carboxylic acid (8.5 mg, 0.027 mmol) and 1,1’-carbonyldiimidazole (6.5 mg, 0.040 mmol) in DMF was stirred at rt for 1hr. After cooling down to 0°C, tert-butyl hydrazinecarboxylate (10.8 mg, 0.081 mmol) in THF (0.6 mL) was added to the stirred mixture. After being stirred at ambient temperature for 16 hrs, the reaction mixture was partitioned between DCM and water. Organic layer was separated and concentrated. The residue was purified by silica gel column chromatography (0 to 10% MeOH in DCM) to give the title compound (11 mg, 75%) as a colorless gum. 1H NMR (500 MHz, Methanol-d4) δ 8.29 (d, J = 7.4, 1H), 8.28 (S, 1H), 7.97 (d, J = 7.4 Hz, 1H), 2.85 – 2.71 (m, 2H), 2.71 – 2.60 (m, 1H), 2.22 – 2.12 (m, 1H), 2.05 (s, 3H), 1.52 (s, 3H).

#### ((*R*)-2-(3-methyl-2,6-dioxopiperidin-3-yl)-5-(5-thioxo-4,5-dihydro-1,3,4-oxadiazol-2-yl)isoindoline-1,3-dione)) (TXM-02-118)

To the stirred solution of tert-butyl (*R*)-2-(2-(3-methyl-2,6-dioxopiperidin-3-yl)-1,3-dioxoisoindoline-5-carbonyl)hydrazine-1-carboxylate (11 mg) in DCM (0.2 mL) was added TFA (0.1 mL) at rt. After being stirred at rt for 1 hr, volatiles were removed under reduced pressure to give a crude product. To the stirred solution of crude (*R*)-2-(3-methyl-2,6-dioxopiperidin-3-yl)-1,3-dioxoisoindoline-5-carbohydrazide and 1,1’-thiocarbonyldiimidazole (7.2 mg, 0.038 mmol) in DMF (0.2 mL) was added TEA (31.2 mL, 0.22 mmol) at rt. After being stirred at 80°C for 16 hrs, the reaction mixture was purified by reverse phase preparative HPLC (0-80% MeOH in water (0.035% TFA modifier)) to give the title compound (1.3 mg) as a pale yellow solid. 1H NMR (500 MHz, DMSO-d6) δ 11.06 (s, 1H), 8.30 (dd, J = 7.8, 1.5 Hz, 1H), 8.14 (d, J = 1.5 Hz, 1H), 8.03 (d, J = 7.8 Hz, 1H), 2.70 (ddd, J = 18.4, 10.5, 5.4 Hz, 1H), 2.65 – 2.54 (m, 2H), 2.13 – 2.04 (m, 1H), 1.92 (s, 3H). RT=0.74 min, m/z: 373.3 [M+H]+.

## Author Contributions

K.C.A., T.H., and S.D.-P. initiated the project. T.M. designed and synthesized reported compounds. H.-S.S. and S.D.-P. carried out crystallographic studies and ITC and DSF experiments. T.H. performed cell experiments. S.D.-P. wrote the manuscript with input from all authors.

## Acknowledgments

We thank Dennis Buckley for engaging discussions, Steve DeAngelo for performing bacterial expressions, and Ezekiel Geffken for performing TR-FRET experiments. S.D.P acknowledges funding from the Linde Family Foundation, the Doris Duke Charitable Foundation, Deerfield 3DC, and Taiho Pharmaceuticals. This work was based upon research conducted at the Northeastern Collaborative Access Team beamlines, which were funded by the National Institute of General Medical Sciences from the National Institutes of Health (P30 GM124165). The Eiger 16M detector on the 24-ID-E beam line was funded by a NIH-ORIP HEI grant (S10OD021527). This research used resources of the Advanced Photon Source, a U.S. Department of Energy (DOE) Office of Science User Facility operated for the DOE Office of Science by Argonne National Laboratory under Contract No. DE-AC02-06CH11357.

## Notes

### Competing Interest Statement

The authors have declared no competing interest.

